# Dual regulation of spine-specific and synapse-to-nucleus signaling by PKCδ during plasticity

**DOI:** 10.1101/2021.09.17.460844

**Authors:** Lesley A. Colgan, Paula Parra-Bueno, Heather L. Holman, Mariah F. Calubag, Jaime A. Misler, Xun Tu, Ryohei Yasuda

## Abstract

The activity-dependent plasticity of synapses is believed to be the cellular basis of learning. These synaptic changes are mediated through the coordination of local biochemical reactions in synapses and changes in gene transcription in the nucleus to modulate neuronal circuits and behavior. The protein kinase C (PKC) family of isozymes has long been established as critical for synaptic plasticity. However, due to a lack of suitable isozyme-specific tools, the role of the novel subfamily of PKC isozymes is largely unknown. Here, through the development of FLIM-FRET activity sensors, we investigate novel PKC isozymes in synaptic plasticity in mouse CA1 pyramidal neurons. We find that PKCδ is activated downstream of TrkB and that the spatiotemporal nature of its activation depends on the plasticity stimulation. In response to single spine plasticity, PKCδ is activated primarily in the stimulated spine and is required for local expression of plasticity. However, in response to multi-spine stimulation, a long-lasting and spreading activation of PKCδ scales with the number of spines stimulated and, by regulating CREB activity, couples spine plasticity to transcription in the nucleus. Thus, PKCδ plays a dual functional role in facilitating synaptic plasticity.

## Introduction

Learning induces structural and functional changes in subsets of active spines to modulate neuronal circuits and behavior (Abdou et al., 2018; Gobbo & Cattaneo, 2020; Hayashi-Takagi et al., 2015). These changes require early local plasticity of spines that is stabilized at later time points through gene transcription in the nucleus. The complex signaling cascades that mediate these changes, therefore, must transduce local and transient inputs (ηm and ms) into long-lasting changes in cell-wide protein expression (mm and hours). The coordination of early local events at spine heads and later changes that occur in the nucleus is not well understood, but kinase cascades, which can extend local, transient signals in both space and time have been implicated (Smolen et al., 2019).

One of the first kinase families identified to be essential for spine plasticity, learning and memory was the protein kinase C (PKC) family (Farley & Auerbach, 1986; Hu et al., 1987; Malenka et al., 1986; Olds et al., 1989). Consisting of more than 12 isozymes and grouped in three subfamilies, it is increasingly clear that different PKC isozymes play unique roles in plasticity. Even single isoforms, whose specificity of signaling is largely defined by their spatiotemporal patterns of activation (Mukherjee et al., 2016), likely have complex roles depending on the nature of the plasticity-inducing stimulus. This complexity, combined with limited existing tools to measure and distinguish PKC isozyme activity with high spatiotemporal resolution, has limited understanding of the processes by which plasticity is expressed and stabilized at the molecular level.

Recently, tools to study the classic subfamily of PKC isozymes in plasticity were developed to reveal an isozyme-specific role for PKCα in the plasticity of spines and heterosynaptic plasticity (Colgan et al., 2018). This isozyme was activated for seconds in single dendritic spines but was able to integrate signals with variable spatiotemporal domains to facilitate metaplasticity and learning. The atypical isozymes, on the other hand, such as PKMζ have been proposed to regulate the maintenance of long-lasting forms of memory on the timescale of hours to days, although their requirement and mechanisms in memory maintenance remain unclear (Borodinova et al., 2017). The role of the novel subfamily of PKC isozymes in synaptic plasticity, however, remains largely unknown.

The novel PKC isozymes consist of four isozymes, three of which are expressed in the mammalian hippocampus (PKCδ, PKCε, PKCη, Naik et al., 2000). Unlike the classic subfamily, novel PKC isozymes do not bind calcium and are primarily activated by binding lipids, particularly the diacylglycerol (DAG) lipid messenger (Steinberg et al., 2008). Here, through loss of function studies and the use of newly-developed, isozyme-specific sensors, we explore the functions of novel PKC isozymes (PKCδ, PKCε, PKCη) in plasticity. We find that PKCδ is uniquely required for the induction of structural plasticity of dendritic spines. Further, we identify its activators and characterize a dual functional role of PKCδ in both local induction of spine plasticity and the stabilization of this plasticity through spine-to-nucleus signaling.

## Results

### PKCδ, but not PKCε or PKCη activity, is required for excitatory synaptic potentiation

To assess the potential role of novel PKC isozymes in synaptic plasticity, we performed loss of function studies in a robust model of synaptic potentiation and learning: structural plasticity (sLTP) of dendritic spines (Choi et al., 2018; Hayashi-Takagi et al., 2015; Matsuzaki et al., 2004). sLTP was induced through optical uncaging of glutamate (30 pulses at 0.5 Hz) in the presence of tetrodotoxin in a Mg^+2^-free aCSF. This allowed for the precise and robust induction of sLTP at a single dendritic spine of CA1 pyramidal neurons in the stratum radiatum of organotypic hippocampal brain slice. sLTP was measured as the change in volume of the stimulated spine. Consistent with previous literature (Harvey, Yasuda, et al., 2008; Matsuzaki et al., 2004), glutamate uncaging induced rapid growth in stimulated spines of wild-type mice. The growth decayed slightly over minutes but remained elevated compared to prestimulus spine size. These changes were both spine-specific and long-lasting. Spine plasticity was not impaired in CA1 neurons in slices prepared from PKCε or PKCη knockout (KO) mice (Figure 1A, B). However, PKCδ KO mice showed impaired induction and expression of spine plasticity (Figure 1C, D). Spine growth was reduced in the early phases of sLTP and by 30 min the volume of the stimulated spine returned to its prestimulus size. This deficit could be fully rescued by sparse, acute (∼24-48 h) overexpression of PKCδ-GFP, indicating the deficit is due to a postsynaptic, cellular role of PKCδ and not due to developmental or circuit-level changes (Figure 1).

**Figure 1:**
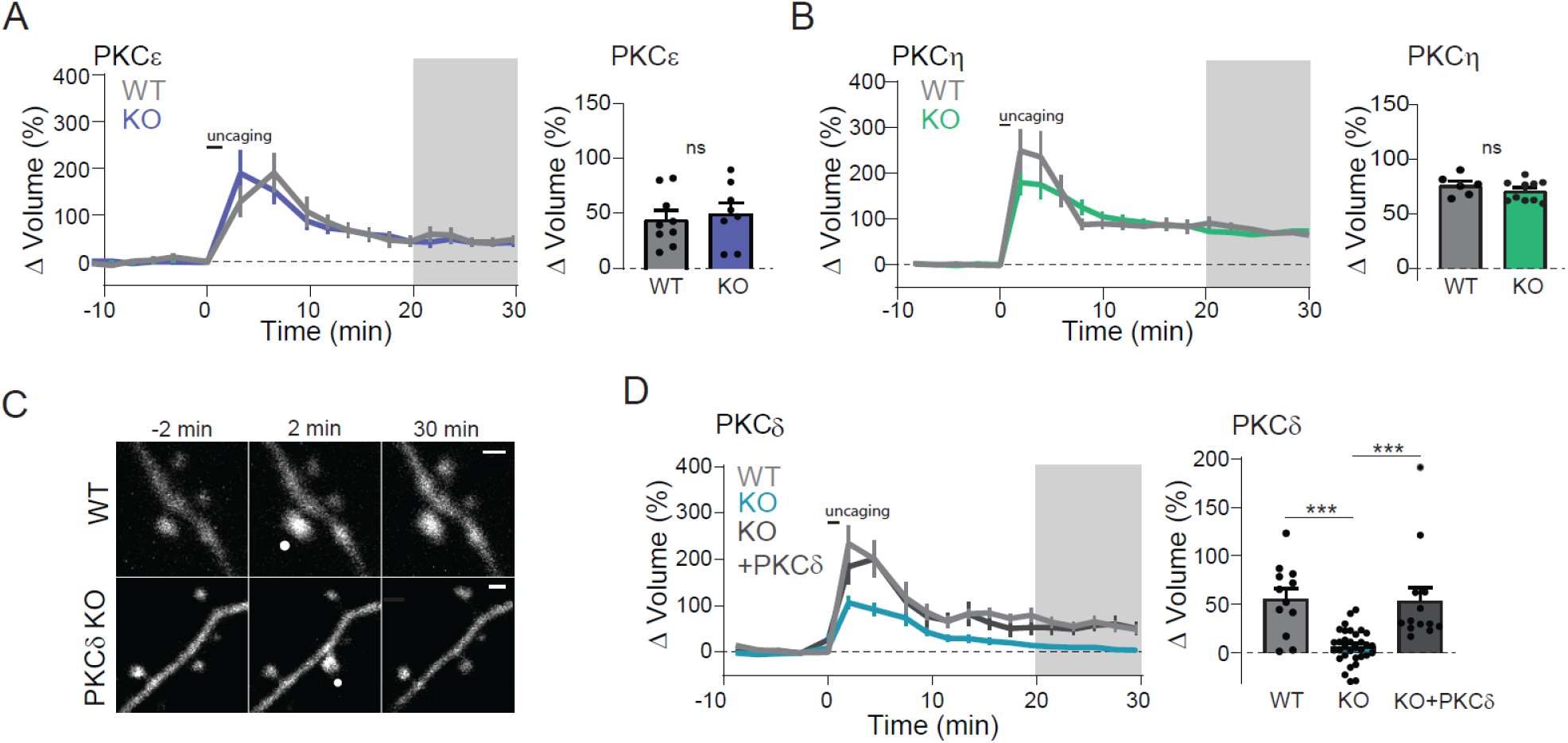
PKCδ, but not PKCε or PKCη, is required for structural plasticity. A, B) Time course and quantification of glutamate uncaging induced change in volume of stimulated dendritic spines in hippocampal CA1 neurons from PKCε WT (n(spines/neurons = 9/6) or KO (n = 8/6) littermates (A) and PKCη WT (n = 6/5) or KO (n = 10/8) littermates (B). Gray shading indicates the time of quantification of mean volume change (right). Two-way, unpaired t-test non-significant ns p > 0.37. C) Representative images of uncaging induced sLTP in neurons from PKCδ WT or KO littermates. The white dot indicates the location of uncaging. Scale bar = 1μm. D) Time course and quantification of change in volume of stimulated spines from PKCδ WT (n = 12/6), KO (n = 33/15), and KO neurons acutely overexpressing GFP-tagged PKCδ (n = 13/7). One-way ANOVA [F (2, 55) = 16.85] p<0.0001. Asterisks indicate Sidak’s post-test comparing indicated groups, *** p<0.001.

As sLTP has been shown to correlate strongly with functional plasticity (Matsuzaki et al., 2004), our results suggested that this structural deficit would be concomitant with deficits in functional, electrically-induced potentiation. To test this, electrophysiological long-term potentiation was induced in acute hippocampal slices made from PKCδ KO or WT littermate animals. Field EPSPs were recorded in the stratum radiatum of the CA1 region before and after stimulation of Schaffer collaterals with a theta-burst stimulation protocol. While no significant differences were seen in basal synaptic transmission (Figure 2A), neurons lacking PKCδ had significantly impaired potentiation (Figure 2B, 2C).

**Figure 2:**
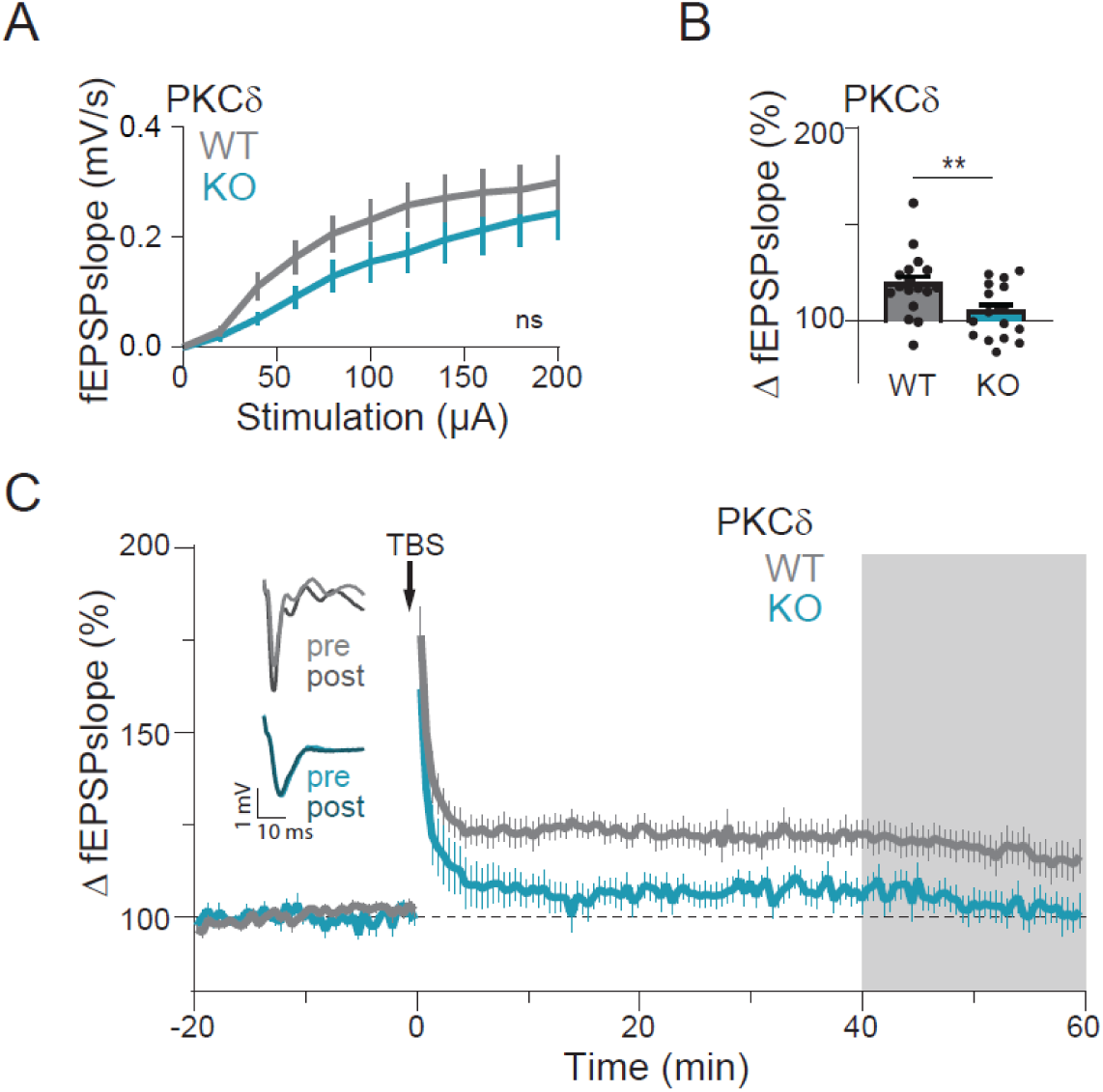
PKCδ is required for functional plasticity. A) Input-output curve of the field EPSP slopes between PKCδ WT and KO littermate mice. ns indicates results of repeated measures Two-Way Anova by genotype [F (1, 30) = 1.829, p=0.19] B, C) Quantification (B) and time course (C) of change in field EPSPs’ slope in PKCδ WT (n = 17/8) and PKCδ KO (n = 16/4) CA1 hippocampal neurons after field stimulation of Schaffer collaterals with a 3x or 5x TBS stimulation protocol. Gray shading indicates the time of quantification in B. Insets (left) are the representative traces of field EPSP’s before (pre) and after (post) stimulation from WT and KO slices. Asterisks indicate unpaired t-test, **p= 0.009.

### Development of isozyme specific sensors for novel PKC isoforms

As this data demonstrated a critical role of PKCδ in structural and functional plasticity, we sought to investigate the spatial and temporal activity of PKCδ during plasticity. Therefore, we developed and characterized isozyme-specific sensors for PKCδ and the other novel PKC isozymes (PKCε and PKCη). To do this, we extended our previously developed and broadly applicable sensor design for the classic PKC isozymes (Colgan et al., 2018). This approach measures two aspects of PKC activation: (1) translocation of the kinase to the plasma membrane and (2) docking of a pseudosubstrate to the active kinase. These two approaches are named ITRACK (**I**sozyme-specific **Tra**nslocation of **C K**inase, Figure 3A top) and IDOCKS (**I**sozyme-specific **Do**cking of **C K**inase **S**ubstrate, Figure 3A bottom) respectively. ITRACK consists of the novel PKC of interest, N-terminally tagged with a FRET donor fluorophore, mEGFP, and expressed with a plasma membrane-targeted FRET acceptor fluorophore, mCherry. Upon activation of the PKC isozyme, translocation of the isozyme to membrane results in FRET between the donor and acceptor fluorophores. This increase in FRET is detected as a decrease in the fluorescence lifetime of the mEGFP-tagged PKC isozyme. IDOCKS similarly consists of an mEGFP-tagged PKC isozyme expressed with an acceptor construct that consists of a pseudosubstrate sequence tagged with two mCherry fluorophores. Upon activation, the acceptor construct binds the active PKC isozyme, increasing FRET between the GFP-tagged novel PKC and the acceptor construct. The activation of the novel PKC of interest, therefore, can be detected as a decrease in the fluorescence lifetime of the donor fluorophore. These designs provide a robust, isozyme-specific measurement of novel PKC activity.

**Figure 3:**
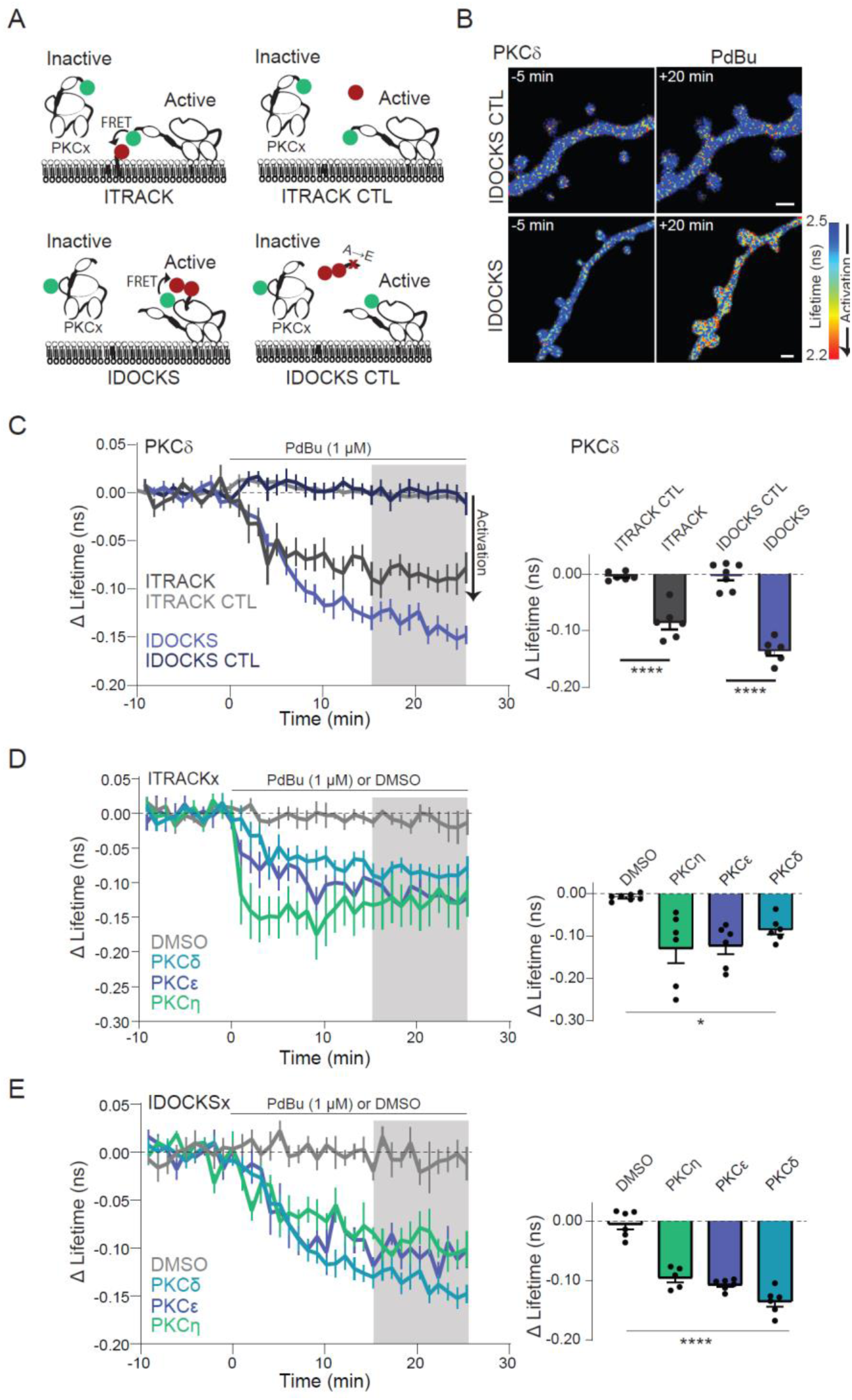
Characterization of FLIM-FRET sensors for novel PKC isozymes in neurons. A) Schematic of **I**sozyme-specific **TRA**nslocation of **C**-**K**inase (ITRACK) and **I**sozyme-specific **DO**cking of **C**-**K**inase **S**ubstrate (IDOCKS) sensors and respective control sensors (ITRACK CTL, IDOCKS CTL). B) Representative lifetime images of IDOCKS and IDOCKS CTL for PKCδ in non-transgenic hippocampal CA1 neurons in response to PKC activation by phorbol ester (1μM PdBu). Warmer colors indicate lifetime decrease and sensor activation. C) Time course and quantification of mean change in the lifetime of ITRACK, IDOCKS, and CTL sensors for PKCδ in response to PdBu. Gray shading indicates the time over which average change was quantified (right). One Way ANOVA [F (3, 21) = 57.85] p<0.0001, Asterisks indicate significance between indicated comparisons tested by Sidak’s multiple comparison post-test p< 0.001. n ≥ 6 neurons. D, E) Time course and quantification of the change in the lifetime of ITRACKx (D) or IDOCKSx (E) for PKCδ, PKCε, or PKCη, measured in hippocampal CA1 neurons in response to bath application of PKC activating drug PdBu or DMSO vehicle as indicated. Gray shading indicates the time over which the average change was quantified. One Way ANOVA [F (3, 19) = 54.98] p<0.0001, Asterisks indicate the lowest level of significance between each PdBu and vehicle group tested by Sidak’s multiple comparison post-test. *p<0.05, ****p<0.001, n ≥ 5.

ITRACK and IDOCKS were tested in cultured cells (Figure S1 A-D) and CA1 pyramidal neurons of organotypic hippocampal slices (Figure 3) by application of a strong pharmacological PKC activator (PdBu, 1 μM). For each of the novel isozymes, both ITRACK and IDOCKS showed significant changes in fluorescence lifetime upon phorbol ester application compared to vehicle application (Figure 3D, E). Importantly, control sensors, in which the acceptor was no longer targeted to the plasma membrane (ITRACK_CTL_) or in which the pseudosubstrate contained a single point mutation that disrupts binding to the PKC kinase site (IDOCKS_CTL_), showed no significant lifetime changes in response to phorbol ester (Figure 3B, C, S1B). This suggests that the lifetime changes observed by ITRACK and IDOCKS reflect translocation of the novel PKCs to the membrane and binding of the novel PKCs to the pseudosubstrate, respectively. Our results suggest that these sensor approaches enable dynamic measurements of novel PKC activity in cells and neurons.

### PKCδ is activated in stimulated spines during plasticity induction

Using our newly developed sensors, we measured the spatial and temporal activation pattern of PKCδ during the induction of single spine plasticity. To do this, we introduced IDOCKSδ into hippocampal CA1 neurons of organotypic slices from PKCδ KO animals and monitored sensor activation during uncaging-induced sLTP. During plasticity induction, there was a rapid decrease in the fluorescence lifetime of IDOCKSδ in the stimulated spine and, to a lesser extent, the underlying dendrite (Figure 4A). This activation of PKCδ peaked during uncaging in the stimulated spine (∼1 min) and in the underlying dendrite with a slightly slower time course. Activation decayed by about 70% over the next several mins but did not return completely to baseline within the time imaged (Figure 4B). Importantly, control stimulations, in the absence of glutamate, or when NMDA receptors were blocked, did not induce activation of PKCδ (Figure 4C), suggesting that activation is induced downstream of NMDA receptors and is not an artifact of laser stimulation. Moreover, the volume change induced during sLTP was not significantly different in neurons expressing either of the two sensors (ITRACKδ or IDOCKSδ) or GFP-tagged PKCδ, suggesting that sensor expression did not disrupt downstream PKCδ signaling (Figure S2A). Unlike the activation seen through IDOCKSδ, ITRACKδ, which measures translocation of PKCδ to the plasma membrane, was not activated during sLTP (Figure S2B, C). This was despite robust activation of ITRACKδ in response to PdBu application in neurons (Figure 3D). This finding suggests that PKCδ is not translocating to KRas-CAAX targeted membrane structures during the induction of sLTP and suggests that the nature and location of PKC isozyme activation are highly dependent on the specific stimulation received.

**Figure 4:**
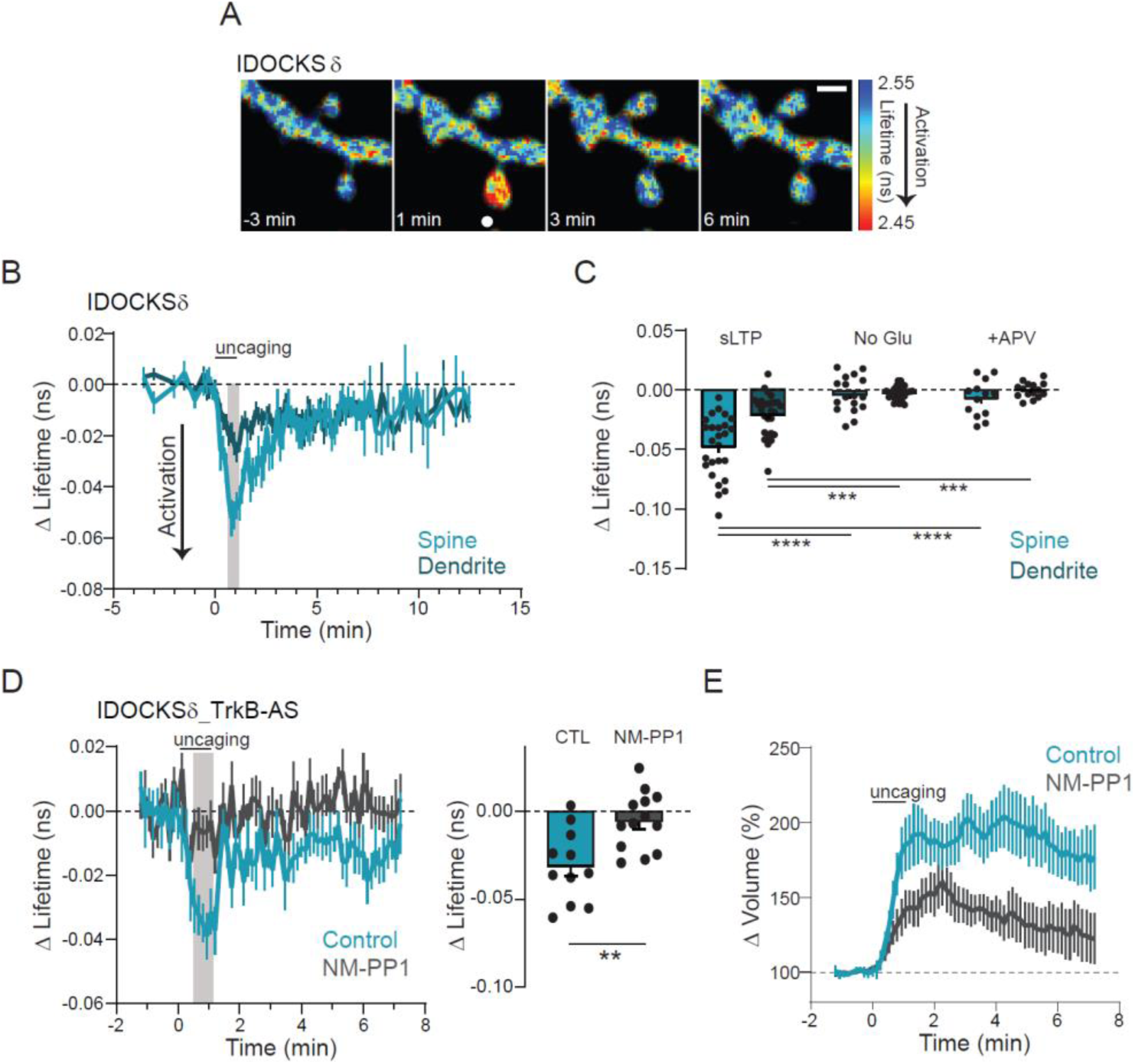
PKCδ is transiently activated in stimulated spines during plasticity. A) Fluorescence lifetime images of IDOCKS in a single spine undergoing sLTP. The white dot indicates uncaging location. Scale bar = 1 μm. Warmer colors indicate shorter lifetimes and PKCδ activation. B) Mean time course of PKCδ activity in the stimulated spine and underlying dendrite measured by the change in the lifetime of IDOCKS during sLTP. n(spines/neurons) = 26/10. C) Quantification of mean lifetime change at the time indicated by gray shading in (B) and in response to uncaging stimulation in the absence of caged glutamate (No Glu n= 17/9) or the presence of the NMDA-receptor antagonist APV (50 μM, n=13/6). Two way ANOVA significant by treatment [F (2, 105) = 32.96, p<0.0001], Asterisks indicate results of Sidak’s multiple comparisons as indicated (****p<0.0001, ***p=0.0004) D) The mean time course (left) and quantification (right) of PKCδ activity in stimulated spines in neurons containing an inert analog-sensitive mutation in TrkB-AS in the presence of vehicle (CTL, n= 11/5) or inhibitory ATP analog (NM-PP1, 1μM n= 13/5). E) Time course of volume change of stimulated spines in D.

### PKCδ is downstream of TrkB activation

Novel PKC isozymes are primarily activated by the production and binding to the lipid second messenger DAG. Recently, increased DAG production during NMDA stimulation or the induction of sLTP was demonstrated (Codazzi, 2006; Colgan et al., 2018). While there could be multiple sources of stimulation-induced DAG production, one mechanism described to support spine plasticity has been through NMDA-dependent autocrine release of BDNF and subsequent activation of TrkB (Harward et al., 2016). As we already demonstrated the requirement of the NMDAR for PKCδ activation (Figure 4C), we tested if PKCδ was also downstream of TrkB activation. IDOCKSδ was therefore monitored in neurons that had been genetically modified to contain an analog-sensitive point mutation in TrkB (TrkB-AS). This mutation is a single-point mutation in the ATP binding pocket of TrkB. The mutation leaves its kinase activity unaltered, but renders it sensitive to inhibition by a synthetic ATP analog, NM-PP1 (Chen et al., 2005). In neurons containing the TrkB-AS mutation that were untreated, or treated only with vehicle, sLTP-induced spine growth and PKCδ activation were normal (Figure 4D, E, CTL). However, TrkB-AS containing neurons treated with the inhibitor NM-PP1 showed impaired PKCδ activation and spine volume change during sLTP (Figure 4D, E, NM-PP1). This demonstrates that TrkB is upstream of PKCδ activity during plasticity, suggesting that PKCδ can monitor levels of BDNF-mediated activation of TrkB to facilitate plasticity.

### PKCδ activity is not restricted to stimulated spine

During plasticity induction, the activation of PKCδ was highest in the stimulated spine, but was present at a lower level throughout the small region of dendrite imaged (Figure 4B), suggesting that PKCδ activity might be able to spread some distance from the synapse. Consistently, TrkB activation during plasticity has previously been reported to regulate long-distance signaling in neurons, including signals from synapses to the nucleus (Esvald et al., 2020; Harward et al., 2016; Moya-Alvarado & Bronfman, 2020). Therefore, we hypothesized that TrkB activation of PKCδ might be a mechanism to extend local, transient signals in both space and time. To investigate whether PKCδ activity was able to spread over long distances, IDOCKS was monitored in the primary apical dendrite before and after inducing plasticity in a single dendritic spine or multiple spines spread across the dendritic tree (Figure 5A). The induction of plasticity in multiple spines was previously shown to induce long-distance signaling to the nucleus of another kinase, the extracellular signal-regulated kinase (ERK) (Zhai et al., 2013). In response to plasticity in one spine, there was little activation of PKCδ in the primary dendrite. However, multiple spine stimulation (sequential induction of sLTP in 5 spines on 3 dendrites) led to a significant increase in PKCδ activity in the primary dendrite that was long-lasting (Figure 5B, C). To monitor the integration of multiple spine stimulation, PKCδ activity was imaged in the primary dendrite during the sequential stimulation of 3 spines on different branches with longer time intervals between spine stimulations (∼8 min). In response to this spaced stimulation of 3 spines, PKCδ activity integrated the stimuli in the primary dendrite to an intermediate level between the activation of 1 and 5 spines (Figure 5D, E). This finding suggests that in addition to a local role in regulating plasticity in the spine, PKCδ may also play a long-distance signaling role in response to multiple spine plasticity.

**Figure 5:**
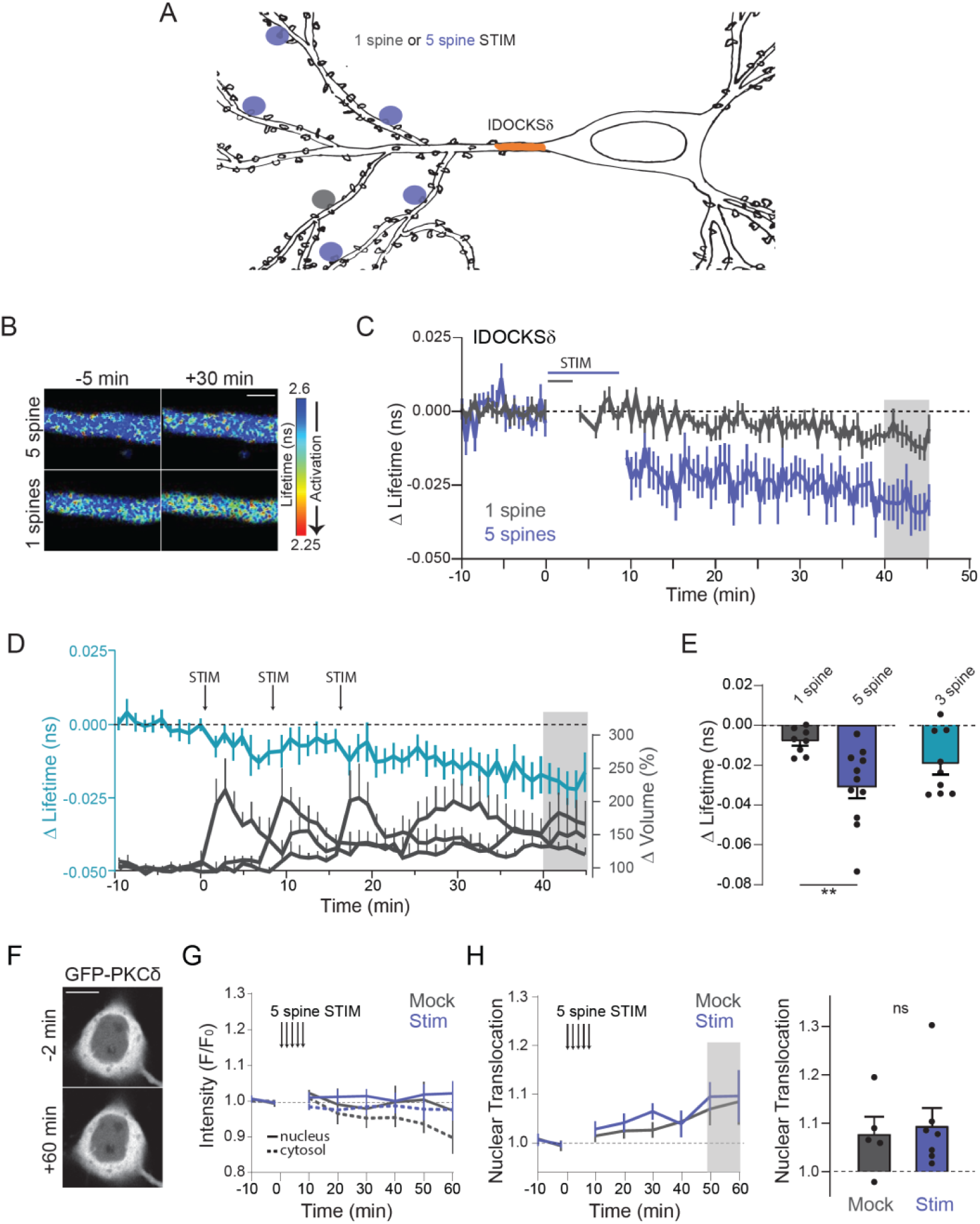
Multispine stimulation enhances long-lasting and spreading activation of PKCδ. A) Schematic of 1 spine or 5 spine stimulation protocol. The gray area indicates the region where PKCδ activation was measured (primary dendrite). B) Representative lifetime images of IDOCKSδ in the primary dendrite in response to 1 or 5 spine stimulation. Warmer colors indicate decreased lifetime and activation of PKCδ. Scale bar = 2 μm. C) Time course of PKCδ activity measured by the change in the lifetime of IDOCKS in response to induction of single spine sLTP (n (neurons) = 8) or sequential induction of sLTP in 5 spines (n=13). D) Time course of PKCδ activation in the primary dendrite (left axis) during the spaced induction of 3 spines. The right axis shows the change in the volume of stimulated spines. n(neurons)= 9. E) Quantification of mean change in the lifetime at 40-45 min (gray area in B) for 1 spine, 5 spines, or 3 spines stimulation (n= 9). Asterisks indicate results from a two-tailed unpaired t-test **p=0.0065. F) Intensity images of GFP-PKCδ in hippocampal CA1 neurons before and 60 min after 5 spines stimulation. Scale bar = 10 μm G) Time course of somatic cytosol and nuclear intensity (F/Fo) of GFP-PKCδ before and after 5 spine stimulation (Stim) or stimulation in the absence of caged Glu (Mock). H) Time course of nuclear translocation (Nuc intensity/Cytosol intensity). Gray shading indicates the time of quantification in H. H) Quantification of mean nuclear translocation after Stim or Mock in C. ns indicated results of unpaired two-tailed t-test. p= 0.7520.

To determine if the long-distance spreading of PKCδ activity was due to the directed translocation of PKCδ into the soma or nucleus, we monitored the somatic localization of GFP-tagged PKCδ (Figure 5F-H). Multi-spine stimulation did not induce significant somatic or nuclear translocation of PKCδ compared to mock stimulation and remained largely excluded from the nucleus. Thus, PKCδ does not show activity-dependent translocation to the somatonuclear compartment but rather, is activated over long distances.

### Acute and long-lasting PKCδ signaling have dual functions

To investigate the potential functional role of this long-distance and long-lasting PKCδ activity further, we needed a tool to selectively inhibit long-lasting PKCδ activity. Due to a lack of isozyme-specific PKC inhibitors, we replaced endogenous PKCδ with a previously characterized PKCδ mutant, PKCδ-AS (Kumar et al., 2015). An analogous mutation to TrkB-AS, the PKCδ-AS mutant consists of a point mutation that leaves its kinase activity unaltered but renders it sensitive to inhibition by a synthetic ATP analog, NM-PP1. Neurons expressing PKCδ-AS in a PKCδ KO background and treated with vehicle showed normal sLTP. However, neurons treated with NM-PP1 shortly before the induction of sLTP showed impaired sLTP (−10 min, Figure 6). The return of spine size to pre-stimulus values at 20-30 min was consistent with the PKCδ KO plasticity phenotype and confirmed that PKCδ kinase activity is required for the spine plasticity. To determine whether sustained PKCδ activity was necessary for the maintenance of early phases of sLTP in the spine, we applied NM-PP1 10 min after spine plasticity was induced. Blocking PKCδ kinase activity after plasticity induction, however, did not impair spine structural plasticity (+10 min, Figure 6 B, C). Therefore, unlike the early activity, long-lasting PKCδ activity is not required for the maintenance of early sLTP of dendritic spines.

**Figure 6:**
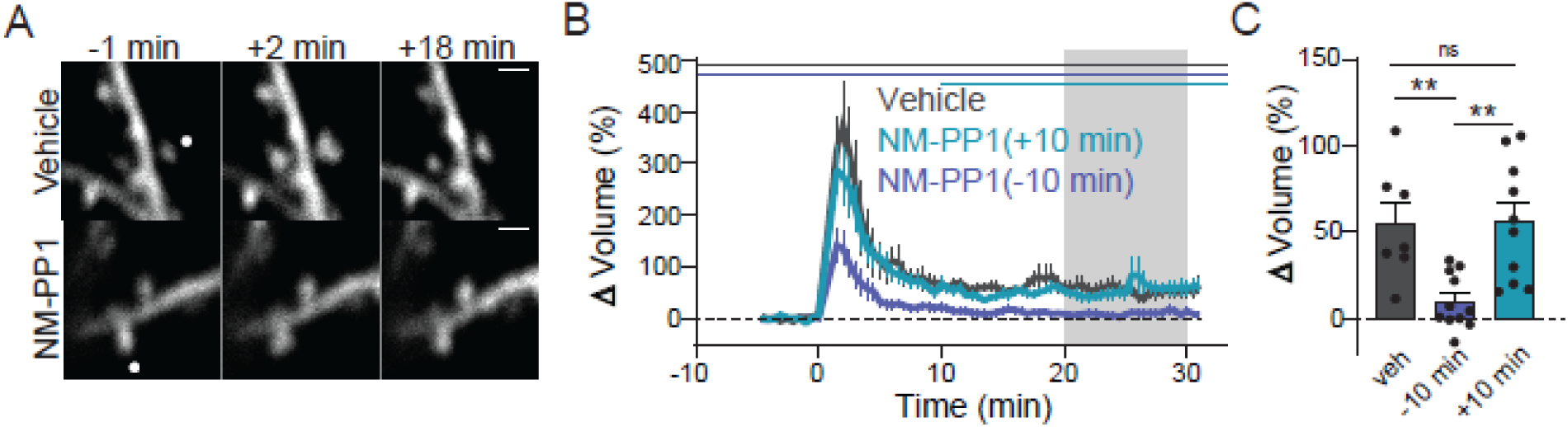
Acute but not long-lasting PKCδ kinase activity is required during plasticity of spines. A) Representative images of sLTP in neurons expressing PKCδ with an inert ATP binding pocket mutation (PKCδ-AS) in place of PKCδ. sLTP was induced in the presence of vehicle or the ATP analog (NM-PP1, 1μM) to inhibit PKCδ kinase activity. The white dot indicates the location of uncaging. scale bar = 1μm. B) Averaged time course of induced volume change of stimulated spines in the presence of vehicle (n=7) or NM-PP1 applied 10 min before (−10 min, n=11) or 10 min after (+10 min, n=10) uncaging stimulation. Gray shading indicates the time of quantification in C. C) Quantification of mean volume change. One way ANOVA [F (2, 25) = 8.688]. Asterisks indicate Sidak’s multiple comparison post-test ** p<0.0089, ns p= 0.999.

### Long-lasting PKCδ activity is required for spine-to-nucleus signaling

The ability of PKCδ to integrate the plasticity of multiple spines into a long-lasting and spreading signal suggested that it may regulate plasticity-dependent transcriptional programs, which are required for late phases of plasticity and memory formation (Alberini & Kandel, 2015). Multi-spine stimulation was previously shown to induce long-lasting activation and nuclear translocation of ERK to regulate spine-to-nucleus signaling (Zhai et al., 2013). Interestingly, this signaling was found to be dependent on long-lasting PKC activity. Therefore, we tested whether ERK was a downstream target of long-lasting PKCδ activity. To do this, PKCδ-AS was expressed in a PKCδ KO background to render it sensitive to inhibition and expressed together with a previously developed FLIM sensor for ERK activity (EKAR, Figure 7A) (Harvey, Ehrhardt, et al., 2008). We directly measured ERK activity in the primary dendrite of neurons during multi-spine stimulation. Consistent with previous data, ERK was activated in a long-lasting manner (Figure 7B, C). However, applying NM-PP1 after the stimulation to inhibit long-lasting PKCδ activity, did not affect ERK activity in the primary dendrite, suggesting that ERK is not a downstream target of this phase of PKCδ activity.

**Figure 7:**
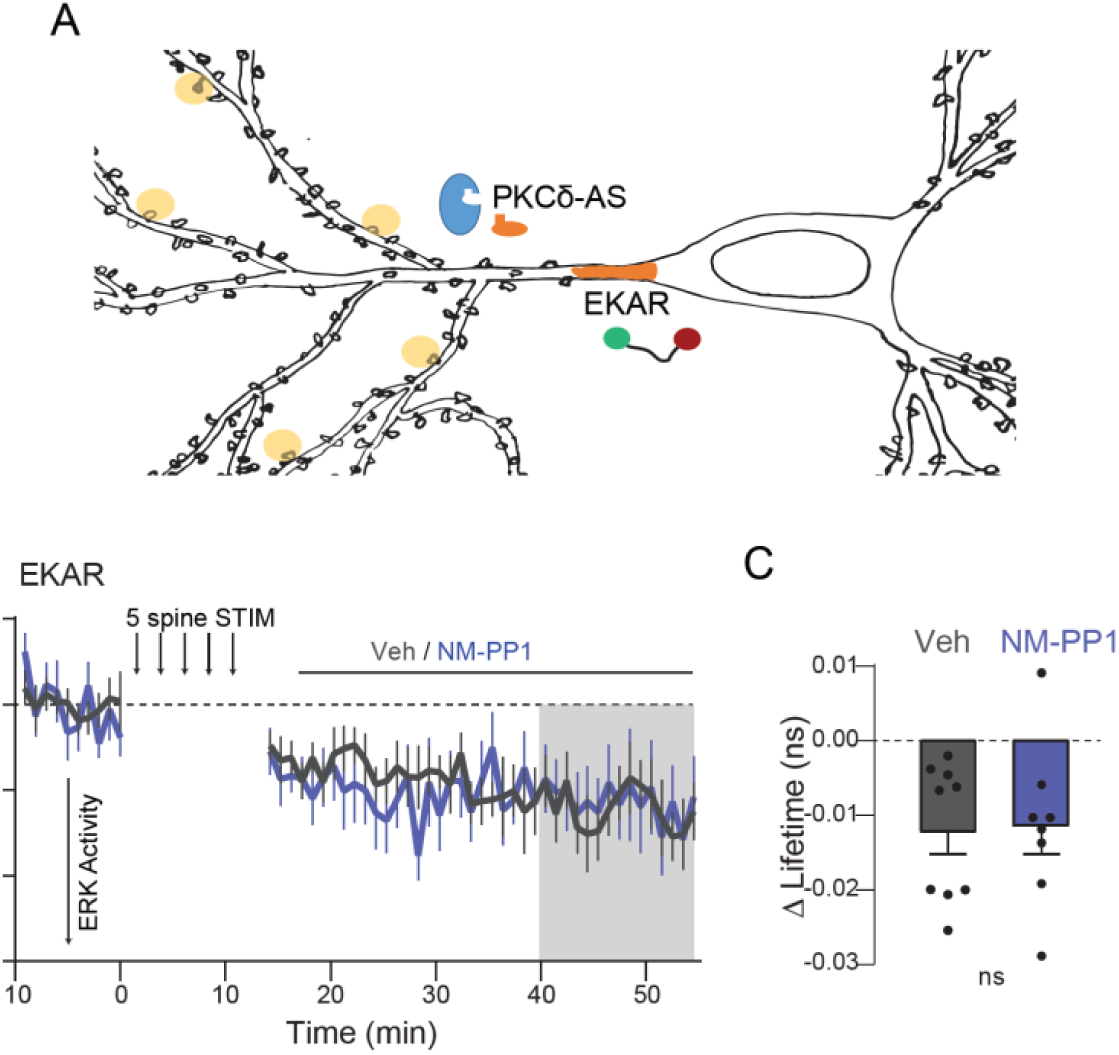
Long-lasting PKCδ activity does not regulate ERK. A) Schematic of experimental design B) The time course of ERK activity as measured by the mean change in the lifetime of the EKAR sensor in the primary dendrite after 5 spine stimulation. ERK activity was compared when either vehicle (n(neurons)= 9) or NM-PP1 (n = 8) was added 10 min after the end of the stimulation to inhibit long-lasting PKCδ activity. C) Quantification of mean change in lifetime at the time indicated by gray shading in B. ns indicates the results of unpaired two-tailed t-test.

We, therefore, tested whether PKCδ might regulate plasticity-induced transcription independent of ERK. The activity-dependent transcription factor cAMP response-element binding protein (CREB) is activated by plasticity-inducing electrical protocols, as well as learning, and is required for the conversion of early to late phases of plasticity (Kandel, 2012). Moreover, PKCδ regulation of CREB has been demonstrated *in-vitro* and in other cellular contexts including in cardiac signaling (Garg et al., 2013; Ozgen et al., 2008; Yamamoto et al., 1988; Zhao, 2007). Therefore, we tested whether long-lasting PKCδ activity is involved in plasticity-induced, synapse-to-nucleus signaling to regulate CREB activity. To do this, PKCδ-AS and a previously developed sensor for CREB were expressed in PKCδ KO neurons and sLTP was sequentially induced in 5 spines (Figure 8A, Laviv et al., 2020). After stimulation, NM-PP1, to inhibit long-lasting PKCδ activity, or vehicle was applied. Multi-spine stimulation led to robust activation of CREB in the nucleus that increased over tens of minutes (Figure 8B, C). However, inhibition of PKCδ with NM-PP1, significantly impaired CREB activation, which plateaued at the time of inhibitor application (Figure 8B-D). This data suggests that PKCδ serves a dual functional role. In addition to facilitating local spine plasticity, PKCδ integrates multi-spine plasticity to regulate the magnitude of CREB activity in the nucleus, an efficient mechanism of spine-to-nucleus signaling.

**Figure 8:**
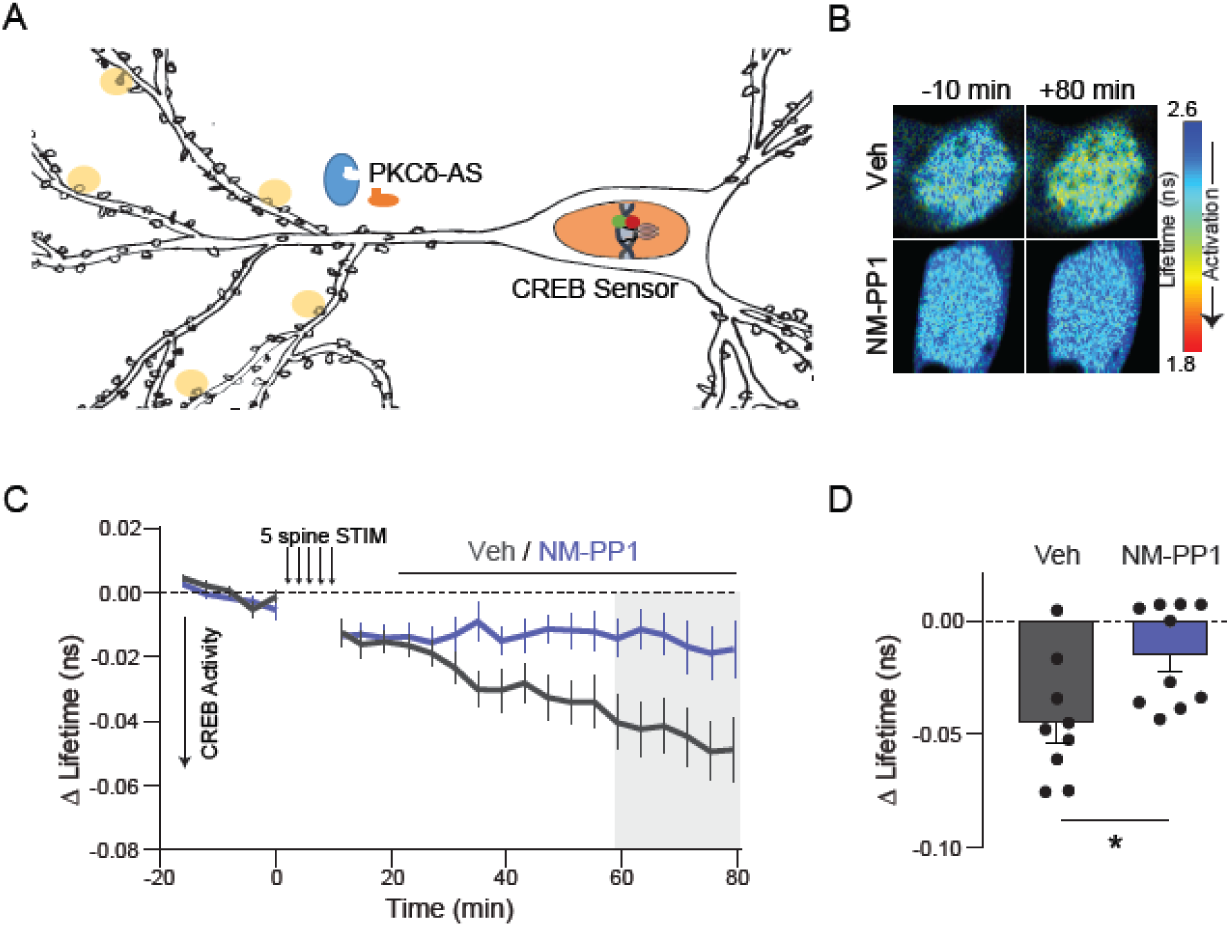
Long-lasting PKCδ activity regulates CREB activity. A) Schematic of five spine stimulation of CA1 neurons in which PKCδ was replaced by PKCδ-AS. CREB activity was monitored in the nucleus using a CREB sensor. B, C, D) Representative lifetime images (B), time course (C) and quantification (D) of mean lifetime change of CREB sensor in the nucleus before after 5 spine stimulation and vehicle (n = 9 neurons) or NM-PP1 (1μM, n = 10 neurons) application 10 min after stimulation to inhibit long-lasting PKCδ activity. Gray shading indicates the time of quantification in D. Asterisk indicates results of two-tailed unpaired t-test *p=0.016

## Discussion

In this study, we have found that PKCδ, amongst the novel isozymes, is uniquely required for the induction of sLTP. Moreover, through the development of highly sensitive isozyme-specific biosensors, we have identified a dual functional role for PKCδ that is defined by the spatiotemporal nature of its activation. During the plasticity of a single spine, NMDA-dependent TrkB activation leads to local PKCδ activity in the stimulated spine within 1 min that returns close to basal levels over approximately 5 min. This early and local activation is essential for the induction of structural and functional spine plasticity. However, the induction of plasticity at multiple spines across the dendritic tree induces a long-lasting (>40 min) PKCδ activation that spreads throughout the neuron. This long-lasting activity scales with the number of stimulated spines to couple plasticity and transcription through the regulation of CREB in the nucleus. This dual functional role of PKCδ in plasticity is an efficient mechanism of coordinating the induction of plasticity in spines with protein synthesis in the nucleus required for long-term plasticity stabilization.

BDNF/TrkB signaling regulates the induction, expression, and consolidation of plasticity as well as learning (Figurov et al., 1996; Kang et al., 1997; Korte et al., 1995; Linnarsson et al., 1997; Minichiello et al., 1999; Mu et al., 1999). One mechanism of this regulation is through the activation of CREB-mediated transcription (Finkbeiner et al., 1997; Ying et al., 2002). This mechanism has often been hypothesized to occur through activation of the Ras/MAPK/ERK signaling (Impey et al., 1998; Ying et al., 2002; Zhai et al., 2013). Our study, however, implicates PKCδ in TrkB signaling to CREB and suggests that its regulation of CREB during plasticity induction is independent of ERK activity. This is consistent with previous work that generated phospho-site specific mutations in TrkB to study the role of specific downstream signaling cascades in plasticity (Minichiello et al., 2002). Animals with a mutation in the phosphosite that binds phospholipase C (PLC) to produce DAG and PKC signaling showed impaired expression and consolidation of LTP and reduced learning (Gruart et al., 2007; Minichiello et al., 2002). Moreover, these neurons showed impaired activation of CREB in response to BDNF application. These deficits were seen although ERK signaling remained intact. Together, the results implicate PKCδ as a necessary mediator of BDNF-TrkB signaling to CREB during CA1 plasticity and hippocampal learning. These results also highlight the need for further understanding of plasticity-dependent transcriptional activation in response to various stimuli. Given the importance of these pathways in both physiology and disease, further study of this diversity of signaling as well as detailed signaling mechanisms through which PKCδ regulates CREB activity is warranted.

The development and characterization of isozyme-specific sensors for novel PKC activity will aid in the further study of the novel class of PKCs. Particularly in investigating the multiple roles of PKC isozymes that are specified by the spatiotemporal nature of their activation. The benefit of multiple sensor designs for protein activation was highlighted in this paper and also serves as a reminder for careful interpretation of sensor results. In this study, PKCδ activation measured by ITRACKδ and IDOCKSδ during sensor characterization showed similar activation profiles in response to a general PKC activating phorbol ester. However, in response to the induction of plasticity via glutamate uncaging, PKCδ activation was seen only with the IDOCKSδ sensor. The ITRACK sensor measures translocation of PKC isozymes to a membrane compartment that is labeled through targeting of a K-ras CAAX domain, which in neurons is primarily plasma membrane (Apolloni et al., 2000). This suggests that during induction of plasticity, PKCδ is active mostly at subcellular compartments that are not labeled by the ITRACKδ acceptor. Consistent with this finding, PKCδ has been shown to translocate to different subcellular domains, including the plasma membrane, the nuclear membrane, and the Golgi, depending on the nature of its activation (Wang et al., 1999), another mechanism to increase its diversity of signaling. Future work to probe PKCδ signaling at particular subcellular compartments, by modifying ITRACKδ to target the acceptor fluorophore to various locations, will help to clarify this diversity of signaling. As the novel subfamily of PKC isoforms has been implicated in numerous disorders, including metabolic disease, cardiovascular disease, autoimmune disorders, and disorders of neurodegeneration, these tools will be of great use for dissecting signaling pathways leading to cellular phenotypes in these diseases.

## Materials and Methods

### DNA constructs

ITRACK: Donor plasmids using restriction site independent cloning were constructed into CMV-promotor containing mEGFP C1 vectors such that mEGFP was fused to the N-terminus of PKCδ (mus musculus), PKCε (mus musculus), and PKCη (mus musculus). The acceptor plasmid, mCh-CAAX, consisted of mCherry followed by a 14 aa linker and a K-Ras derived lipid targeting motif (DGKKKKKKSKTKCVIM) driven by a CMV promoter. Negative controls for ITRACK (ITRACK CTL) consisted of the same donor construct and a control acceptor construct named mCh-CAAXneg in which a stop codon was introduced before the lipid targeting domain. IDOCKS: Donor plasmids consisted of PKCδ, PKCε, and PKCη which were tagged on their C termini with mEGFP driven by CMV promoter. The acceptor construct, 2mCh-PS, consisted of two copies of mCherry fluorophores separated by an eight amino acid linker followed by the 18 amino acid pseudosubstrate region from PKCα/β (RFARKGALRQKNVHEVKN) driven by the CMV promoter. The negative control IDOCKS consisted of the same acceptor construct with a single point mutation R→E (RFARKGAL**E**QKNVHEVKN). GFP-PKCδ-AS was made through a single point mutation M425 to A in the donor construct of GFP tagged PKCδ (Kumar et al., 2015). CREB and EKAR sensors consisted of DNA constructs as previously described (Harvey, Ehrhardt, et al., 2008; Laviv et al., 2020).

### HeLa cell maintenance, transfection and imaging

HeLa cells (ATCC CCL-2) were grown in Dulbecco’s modified Eagle medium supplemented with 10% fetal bovine serum at 37 °C in 5% CO_2_. Plasmids were transfected into HeLa cells using Lipofectamine 2000 (Invitrogen) at a ratio of donor plasmid to acceptor plasmid of 1:3 for ITRACK and 1:2 for IDOCKS. Imaging was performed 24-48h following transfection in a HEPES-buffered aCSF solution (20 mM HEPES pH 7.3, 130 mM NaCl, 2 mM NaHCO_3_, 25 mM D-glucose, 2.5 mM KCl, 1.25 mM NaH_2_PO_4_) with 2 mM CaCl_2_ and 2 mM MgCl_2_ by 2pFLIM as described below. When indicated, cells were stimulated with 1 µM PdBu (Tocris) or DMSO (0.02%) vehicle.

### Nontransgenic and Transgenic Animals

All experimental procedures were approved by the Max Planck Florida Institute for Neuroscience Animal Care and Use Committee, following guidelines by the US National Institutes of Health. Either nontransgenic or transgenic P3-P8 mouse pups from both sexes were used for organotypic slices for imaging studies as indicated. P30-P50 mice of both sexes were used for acute slices for electrophysiological studies.

- Nontransgenic animals, C57Bl/6N Crl, were received from Charles River Lab. These animals were used in Figures 3.
- PKCδ KO mice 129X1/SvJ were obtained from Jaxson labs (JAX stock #028055) and developed as described previously described (Chou et al., 2004). Mice were backcrossed to swiss webster mouse line (CRL CFW) to improve breeding efficiency, litter size and mothering characteristics. Mice were bred het x het to generate WT and KO littermates. These animals were used for Figures 1C,D, 2, 4A-C, 5-8, S2.
- PKCε KO mice B6.129S4-Prkcetm1Msg/J were obtained from Jackson Lab (Jax Stock #004189) and were developed as previously described (Khasar et al., 1999). Mice were bred het x het to generate WT and KO littermates. These animals were used in Figure 1A.
- PKC eta KO mice B6.Cg-Prkchtm1.2Gasc/J were obtained from Jackson Lab (Jax Stock #018988) and were developed as previously described (Fu et al., 2011). Mice were bred het x het to generate WT and KO littermates. These animals were used in Figure 1B.
- Double transgenic mice PKCδKO/TrkB-AS were generated by crossing PKCδKO mice with TrkB-AS mutant mice (TrkB^F616A^ C57bl/6). TrkB^F616A^ C57bl/6 mutant mice were developed and provided by Dr. David Ginty as previously described (Chen et al., 2005). Breeding pairs were KO on one gene and Het on the other gene. These animals were used in Figure 4D, E.

### Organotypic hippocampal slice cultures and transfection

Organotypic hippocampal slices were prepared from non-transgenic or transgenic postnatal 3-8 day old pups of both sexes as previously described (Stoppini et al., 1991). In brief, pups were deeply anesthetized by isofluorane and then euthanized by decapitation. After removing the brain, hippocampi were dissected and cut into 350 µm thick coronal slices using a McIlwain tissue chopper (Ted Pella, Inc). Slices were plated on hydrophilic PTFE membranes (Millicell, Millipore) and maintained at 37degrees and a 5% CO_2_ environment in culture medium (MEM medium (Life Technologies), 20% horse serum, 1mM L-Glutamine, 1mM CaCl_2_, 2mM MgSO_4_, 12.9mM D-Glucose, 5.2mM NaHCO_3_, 30mM Hepes, 0.075% Ascorbic Acid, 1µg/ml Insulin) placed beneath the membranes. The culture medium was exchanged every other day. Organotypic slices were transfected after 7-10 DIV with biolistic gene transfer (O’Brien & Lummis, 2006) using 1.6 µm gold beads (8 mg) coated with plasmids containing DNA of interest in the following amounts. mEGFP: 15 µg, PKCδ_mEGFP: 30 µg, AS PKCδ_mEGFP: 30 µg, ITRACK or ITRACKneg: 15 µg donor (mEGFP-PKCδ, mEGFP-PKCε, or mEGFP-PKCη) and 40 µg acceptor (mCh-CAAX or mCh-CAAX_neg), IDOCKS or IDOCKSneg: 10 µg donor (PKCδ-mEGFP, PKCε-mEGFP, or PKCη-mEGFP) and 20 µg Acceptor (2mCh-PS or 2mCh-PSmut), GFP-PKCδ-AS: 30 μg, GFP-PKCδ-AS and CREB sensor: GFP-AS-PKCδ 15 μg and 15 μg donor (mEGFP-CREB) and 30 μg acceptor (mCherry-KIX-mCherry), PKCδ-GFP: 30 μg, PKCδ-AS and EKAR: 20 μg each. Secondary or tertiary apical dendrites in the stratum radiatum of transfected CA1 pyramidal neurons were imaged 2–7 days after transfection by 2-photon microscopy or 2pFLIM as described below.

### 2-photon microscopy and 2pFLIM

PKC isozyme activity was measured using 2pFLIM. For quantification of spine volume change, we monitored the fluorescence intensity change of mEGFP in the spines using regular two-photon microscopy that was simultaneous with lifetime measurements. Intensity measurements and 2pFLIM imaging in HeLa cells and slices were performed using a custom 2p microscope. mEGFP and mCh were excited with a Ti:sapphire laser (Chameleon, Coherent) at a wavelength of 920 nm and a power of 1.4-1.6 mW measured below the objective. The fluorescence was collected with an objective (60×, 1.0 n.a, Olympus), divided with a dichroic mirror (565 nm), and detected with two separated photoelectron multiplier tubes placed after wavelength filters (Chroma, 510/70-2p for green and 620/90-2p for red). Photoelectron multiplier tubes with low transfer time spread (H7422-40p; Hamamatsu) were used for both red and green channels. The system was controlled via PCIe 6323 and data is acquired by TimeHarp 260 pico dual (Picoquant). Software for data acquisition and analysis is FLIMage (Fluorescence Lifetime Imaging). FLIM images were taken at 64 × 64 or 128 × 128 resolution with 2, 6 or 12 frames averaged. Intensity images for analysis of sLTP volume change were collected by 128×128 pixels as a z stack of three slices with 1 µm separation and averaging 6 frames/slice. Spine volume change was calculated as the background-subtracted F - F_0_ where F_0_ was the average fluorescence intensity before stimulation. In some cases of sLTP volume imaging of PKC KO neurons and WT littermates, parallel automated imaging of 2-3 spines per neuron was done using a custom-built interface in MATLAB by employing algorithms for autofocusing and drift correction to maintain position and focus (Smirnov et al., 2017). For these experiments, a 2 min stagger of uncaging events was incorporated to avoid data loss during uncaging events. Nuclear translocation was calculated from intensity measurements of the nucleus (nuc) and surrounding cytoplasmic fluorescence (cyt) in the central cross-section of the nucleus and calculated as the background-subtracted [F_nuc_/Fo_nuc_]/[F_cyt_/Fo_cyt_], where Fo was the average fluorescence intensity before stimulation.

### Two-photon glutamate uncaging

Structural plasticity of dendritic spines was stimulated through uncaging of 4-Methoxy-7-nitroindolinyl-caged-L-glutamate (MNI-caged glutamate, Tocris) using a Ti: Sapphire laser tuned at a wavelength of 720 nm. The uncaging laser was focused to a small region ∼0.5 µm from the spine and 2.7-2.9mW of laser power (measured at the objective) was pulsed 30 times in a 0.5 Hz train with a 6ms pulse width. Spines deeper than 50 µm were not selected for uncaging experiments. Experiments were performed in Mg^2+^ fee artificial cerebral spinal fluid (ACSF; 127 mM NaCl, 2.5 mM KCl, 4 mM CaCl_2_, 25 mM NaHCO_3_, 1.25 mM NaH_2_PO_4_ and 25 mM glucose) containing 1 µM tetrodotoxin (TTX) and 4 mM MNI-caged L-glutamate aerated with 95% O_2_ and 5% CO_2._ Experiments were performed at room temperature (24-26°C). For multi-spine stimulation experiments, 5 spines were stimulated sequentially on 3-4 secondary or tertiary dendrites. Spines stimulated on the same dendritic branches were spaced by more than 25 μm.

### 2pFLIM analysis

To measure the change in fluorescence lifetime we fit a fluorescence lifetime curve summing all pixels over a whole image with a double exponential function convolved with the Gaussian pulse response function:

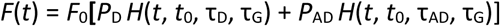

where τ_AD_ is the fluorescence lifetime of the donor bound with the acceptor, *P*_D_ and *P*_AD_ are the fraction of free donor and the donor undergoing FRET with the acceptor, respectively, and *H*(*t*) is a fluorescence lifetime curve with a single exponential function convolved with the Gaussian pulse response function:

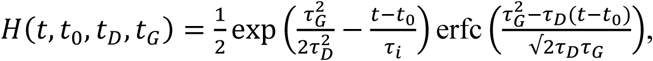

in which τ_D_ is the fluorescence lifetime of the free donor, τ_G_ is the width of the Gaussian pulse response function, *F*_0_ is the peak fluorescence before convolution and *t*_0_ is the time offset, and erfc is the complementary error function.

We fixed τ_D_ to 2.65 ns and τ_AD_ to 1.33 ns based on previously published work of classic PKC isozyme sensors to obtain stable fitting (Colgan et al., 2018). To generate the fluorescence lifetime image, we calculated the mean photon arrival time, <*t*>, in each pixel as:

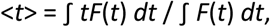

Then the mean fluorescence lifetime, <*τ*>, is calculated as the mean photon arrival time minus an offset arrival time, *t*_*o*_, which is obtained by fitting the whole image:

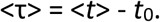

Change in lifetime is calculated as mean fluorescence lifetime <*τ*> in ROI subtracted by the average lifetime in the ROI before stimulation.

The source code of the software is available online: https://github.com/ryoheiyasuda/FLIMage_public

### Pharmacology

1NM-PP1 (Santa Cruz, WD 1 µM), Phorbol 12,13-dibutyrate (PdBu, Tocris, WD 1 µM), MNI Caged Glu (Tocris, 4 mM), APV (Sigma, WD 50 µM) were stored as recommended by supplier and diluted into aCSF to the working dilutions (WD) listed.

### Electrophysiology

Acute slice preparation: PKCδ WT or KO littermate mice (P30-P50) were sedated by isoflurane inhalation, and perfused intracardially with a chilled choline chloride solution. Brain was removed and placed in the same choline chloride solution composed of 124 mM Choline Chloride, 2.5 mM KCl, 26 mM NaHCO3, 3.3 mM MgCl2, 1.2 mM NaH2PO4, 10 mM Glucose and 0.5 mM CaCl2, pH 7.4 equilibrated with 95%O2/5%CO2. Coronal slices (300 µm) containing the hippocampus were cut using a vibratome (Leica) and maintained in a submerged chamber at 32 °C for 1h and then at room temperature in oxygenated ACSF. Extracellular Recordings and LTP protocol: Slices were perfused with oxygenated ACSF containing 2 mM CaCl2, 2 mM MgCl2, and 100 µM picrotoxin. One glass electrode (resistance ∼4 MΩ) containing the same ACSF solution was placed in the dendritic layer of CA1 area (∼100–200 µm away from the soma) while stimulating Schaffer Collateral fibers with current square pulses (0.1 ms) using a concentric bipolar stimulation electrode (FHC). The initial slope of the fEPSP was monitored with custom software (MatLab). The stimulation strength was set to ∼50% saturation. A 20 min stable baseline was first recorded before induction of LTP. LTP was induced by applying 3 to 5 trains of TBS stimulation. fEPSPs responses were recorded for an hour after the stimulation protocol. All data were analyzed with an in-house program written with Matlab.

### Experimental Design and Statistical analysis

All values are presented as mean ± SEM unless otherwise noted. The number of independent measurements (n) is indicated in figure legends. For in-vitro studies, neurons were assigned to different groups to randomly interleave control and experimental groups from slices prepared from the same animals. Experiments from control and experimental groups were randomly interleaved during experimentation with the exception that each block of experiments began with one neuron from the control group to ensure the technical success of experiments. Data distribution was assumed to be normal but this was not formally tested. Unpaired two-tailed student’s t-test was used for comparing two independent samples. One-way ANOVA followed by multiple comparison tests was used for comparing more than two independent samples. Two-way ANOVA followed by multiple comparison tests were used to compare grouped data sets. All t-tests and one-way ANOVAs included formal testing for differences in variance (e.g F test or Bartlett’s test to compare variances) and appropriate corrections were made and are indicated in instances where this was relevant. Statistical tests and p values are noted in each figure legend and were computed using GraphPad Prism 9 for Windows (GraphPad Software, La Jolla California USA, www.graphpad.com). For *in-vitro* experiments, data collection and analysis were not performed blind to the conditions of the experiments. Neurons in which there were obvious signs of poor health that developed during the experiment such as beading of processes were excluded before analysis. FLIM data in which there was more than 0.06 ns variation in the baseline (due to low photon number) were excluded from further analysis.

## Acknowledgments

We would like to acknowledge David Kloetzer for lab management, Long Yan for microscopy development and maintenance, Yuki Hayano for technical assistance, Mary Philips for suggestions in figure design and the MPFI ARC, including Minida Dowdy, Elizabeth Garcia, and Amanda Coldwell for animal care and maintenance. This work was supported by National Institutes of Health Grants R35-NS-116804 (RY), R01-MH-080047 (RY), and F32MH101954 (L.A.C.).

---

**Figure S1:**
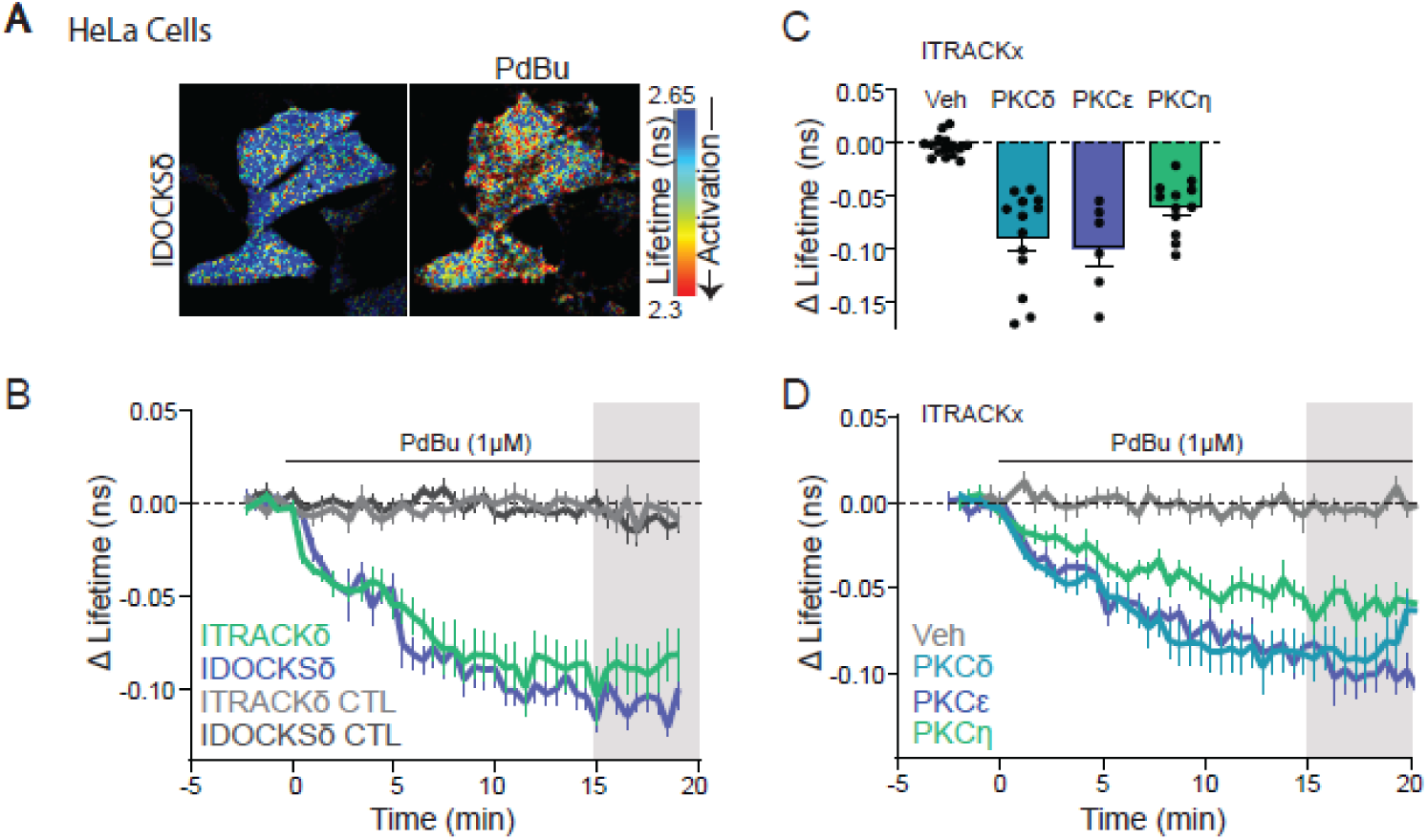
Characterization of novel PKC sensors in HeLa cells. A) Representative lifetime images of IDOCKSδ in HeLa cells before and after PdBU (1 μM) application. Warmer colors indicate a shorter lifetime and activation of PKCδ. B) The time course of mean change in the lifetime of ITRACKδ (n (cells/experiments)= 13/4) and IDOCKSδ (12/3) and control sensors in Hela cells in response to bath application of PdBU (1 μM). C) Quantification of mean lifetime change of PKCδ, PKCε, or PKCη activity measured by ITRACK in response to PdBu or DMSO (Veh) as indicated. D) Time course of the data in C.

**Figure S2:**
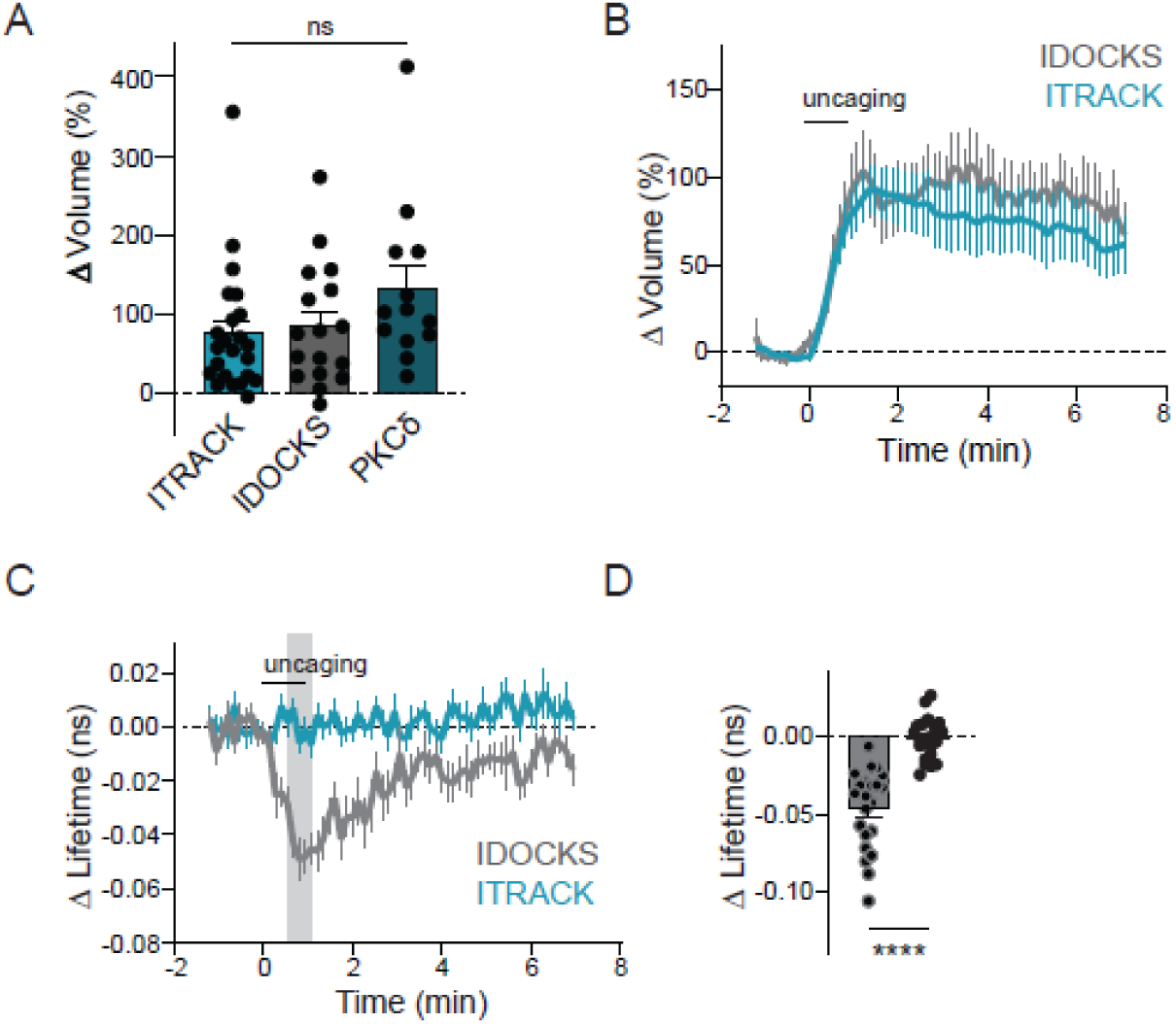
IDOCKSδ but not ITRACKδ show PKCδ activation during sLTP. A) The volume change of stimulated spines of PKCδ KO neurons expressing ITRACKδ, IDOCKSδ, or PKCδ. One way ANOVA ns [F (2, 50) = 1.933] p= 0.16. B) Time course of volume change of stimulated spines in PKCδ KO neurons expressing IDOCKSδ or ITRACKδ. C) Mean time course of PKCδ activity in the stimulated spine measured by the change in the lifetime of IDOCKSδ or ITRACKδ in response to induction of spine plasticity. D) Quantification of the mean change in the lifetime at 0.5 - 1 min (shaded region in C). Asterisks indicate results of an unpaired two-tailed t-test with Welch’s correction for unequal variance ****p<0.0001.

